# DNA methylation is indispensable for leukemia inhibitory factor dependent embryonic stem cells reprogramming

**DOI:** 10.1101/2020.03.17.994939

**Authors:** Baojiang Wu, Yunxia Li, Bojiang Li, Baojing Zhang, Yanqiu Wang, Lin Li, Junpeng Gao, Yuting Fu, Shudong Li, Chen Chen, M. Azim Surani, Fuchou Tang, Xihe Li, Siqin Bao

## Abstract

Naïve pluripotency can be maintained by the 2i/LIF supplements (CHIR99021, PD0325901 and LIF), which primarily affect canonical WNT, FGF/ERK, and JAK/STAT3 signaling. However, whether one of these tripartite supplements alone is sufficient to maintain naïve self-renewal remain unclear. Here we show that LIF alone is sufficient to induce reprogramming of 2i/LIF cultured ESCs (2i/L-ESCs) to ESCs with hypermethylated state (L-ESCs). *In vitro*, upon withdrawal of 2i, 2i/L-ESCs overcome the epigenetic barrier and DNA hypermethylated, which accompanies transcriptional changes and subsequent establishment of epigenetic memory. Global transcriptome features also show that L-ESCs are close to 2i/L-ESCs and in a stable state between naïve and primed pluripotency. Notably, our results demonstrate that DNA methylation was indispensable for LIF-dependent mouse ESCs reprogramming and self-renew. LIF-dependent ESCs reprogramming efficiency is significantly increased in serum treatment and reduced in *Dnmt3a* or *Dnmt3l* knockout ESCs. Importantly, unlike epiblast and EpiSCs, L-ESCs contribute to somatic tissues and germ cells in chimaeras. Such simple culture system of ESCs is more conducive to clarify the molecular mechanism of ESCs *in vitro* culture.

**Significance:** Embryonic stem cell (ESCs) exhibit naïve pluripotency which reflects their ability to contribute to all embryonic lineages upon injection into blastocyst. ESCs were originally derived by co-culture with feeder cells and fetal calf serum. In this manuscript, we took a detailed approach to dissect the roles of LIF alone in ESC reprogramming of 2i/LIF cultured ESCs (2i/L-ESCs). Here, for the first time, we derived stable hypermethylated pluripotent ESCs under culture of LIF alone (L-ESCs). We further assessed L-ESCs properties both in vitro and in vivo, and provide molecular insights to the mechanism which allows LIF alone to maintain pluripotency and a hypermethylated state. We believe these findings are novel and valuable for future ESCs study.

## Introduction

Mouse embryonic stem cells (ESCs) are isolated from the inner cell mass of the pre-implantation embryos (1, 2). Since pluripotent mouse embryonic stem cells were first established four decades ago, various culture systems of ESCs have been developed including initially, using feeder/serum/cytokines, then feeder/serum/Leukemia inhibitory factor (LIF) or Bone morphogenetic protein 4 (BMP4) (3–5), and more recently using 2i/LIF (two inhibitors CHIR99021, PD0325901 and LIF) (6). It is generally believed that the optimal culture condition for ground state ESCs comprises three additive 2i/LIF supplements which affect canonical WNT, FGF/ERK, and JAK/STAT3 signals respectively (7). It has been reported that combination of any two of these tripartite supplements was sufficient to maintain naïve self-renewal of ESCs (8). However, whether any one of the tripartite supplements plays a critical role with unique signalling targets for ESCs pluripotency and self-renewal remains unanswered.

LIF is the most pleiotropic member of the interleukin-6 family of cytokines, and utilizes a receptor that consists of the LIF receptor B and gp130 (7). LIF is able to activate three intracellular signaling pathway: the JAK/STAT pathway, the PI3K/AKT pathway, and the SH2 domain-containing tyrosine phosphatase/MAPK pathway. LIF has antagonistic effects in different cell types including stimulating or inhibiting cell proliferation, differentiation and survival. Since LIF was detected in extract from feeder cells and has been used for most mice ESCs medium, it has been fully demonstrated to be an important supplement for ESCs self-renewal and pluripotency (4–6, 9–12). Nevertheless, essential LIF/STAT3 functions can be compensated by activation of canonical WNT signaling and inhibition of FGF/ERK in the established culture system for self-renewal of ESCs (7). However, the consequences LIF/STAT3 signaling alone and precise regulatory mechanisms for ESCs self-renew have remained largely elusive.

Mouse ESCs cultured in different culture conditions exhibit distinct DNA methylation patterns. The 2i/LIF cultured ESCs (2i/L-ESCs) are globally DNA hypomethylated, whereas ESCs are grown in classical medium containing feeders, serum and LIF (S/L-ESCs) show global DNA hypermethylation (13, 14). Additionally, DNA methylation levels were shown to be reversible between S/L-ESCs and 2i/L-ESCs (13). Recent research reported prolonged MEK1/2 suppression impairs the epigenetic and genomic integrity as well as the developmental potential of ESCs, in part through the downregulation of DNA methylation (15, 16). This suggests that DNA methylation plays an important role in ESCs and normal development. We also showed that hypermethylation is a key point for expanded pluripotency of ESCs in chemical defined medium (17).

The combination of 2i supports the self-renewal of ESCs in serum-free culture without LIF, however, addition of LIF in 2i-culture condition further promoted self-renewal of ESCs, suggesting the synergistic effect of 2i and LIF (6). PD0325901 suppresses the differentiation of ESCs but does not support proliferation (6, 18). CHIR99021 is highly specific to GSK3 and it alone is not sufficient to support the self-renewal of ESCs in serum-free culture (6). In this study, we focus on the JAK/STAT3 signaling and show that LIF alone is able to support mouse embryonic stem cells self-renew and pluripotency as well as developmental potency. Our data also suggest that DNA methylation is indispensable for LIF dependent mouse ESCs reprogramming and self-renew. The detailed analysis of LIF alone dependent mouse ESCs reprogramming provides mechanistic insight into global DNA (de)methylation and also provides a rich resource for future studies on ESCs *in vitro* culture.

## Results

### LIF alone supports ESCs self-renew and pluripotency in chemically defined media

Serum plus LIF (S/L) medium and 2i plus LIF (2i/LIF) medium (based N2B27) are two typical ESCs culture media. In particular, LIF was found in almost all mice ESCs culture media *in vitro*. Therefore, we sought to determine whether LIF alone is capable of driving continuous cycles of self-renew of ESCs in the absence of serum and 2i medium. In here, we used six Oct4-ΔPE-GFP (GOF/GFP, mixed background of MF1, 129/sv, and C57BL/6J strains) ×129/sv F1 mice (19) ESCs lines (W1, W2, W4, W5, W6 and SQ3.3), which were directly derived in 2i/L medium (passages 15-20) and then switched to chemically defined LIF (1000 IU/ml) medium based on N2B27 (L-medium) (Fig. 1A and *SI Appendix*, Fig. S1A). Initially ESCs showed signs of differentiation, such as flattening of colonies and reduction of GOF/GFP positive (GOF/GFP^+^) for pluripotency-related transcription factors *Oct4* (Fig. 1B). However, in passages 3-5, some GOF/GFP^+^ colonies similar to those in undifferentiated ESCs, began to survive during LIF dependent ESCs reprogramming (Fig. 1B). Here we designated these LIF-dependent GOF/GFP^+^ ESCs in chemically defined medium as L-ESCs. GOF/GFP^+^ colonies increased gradually with further passages (Fig. 1B).

**Figure 1.**
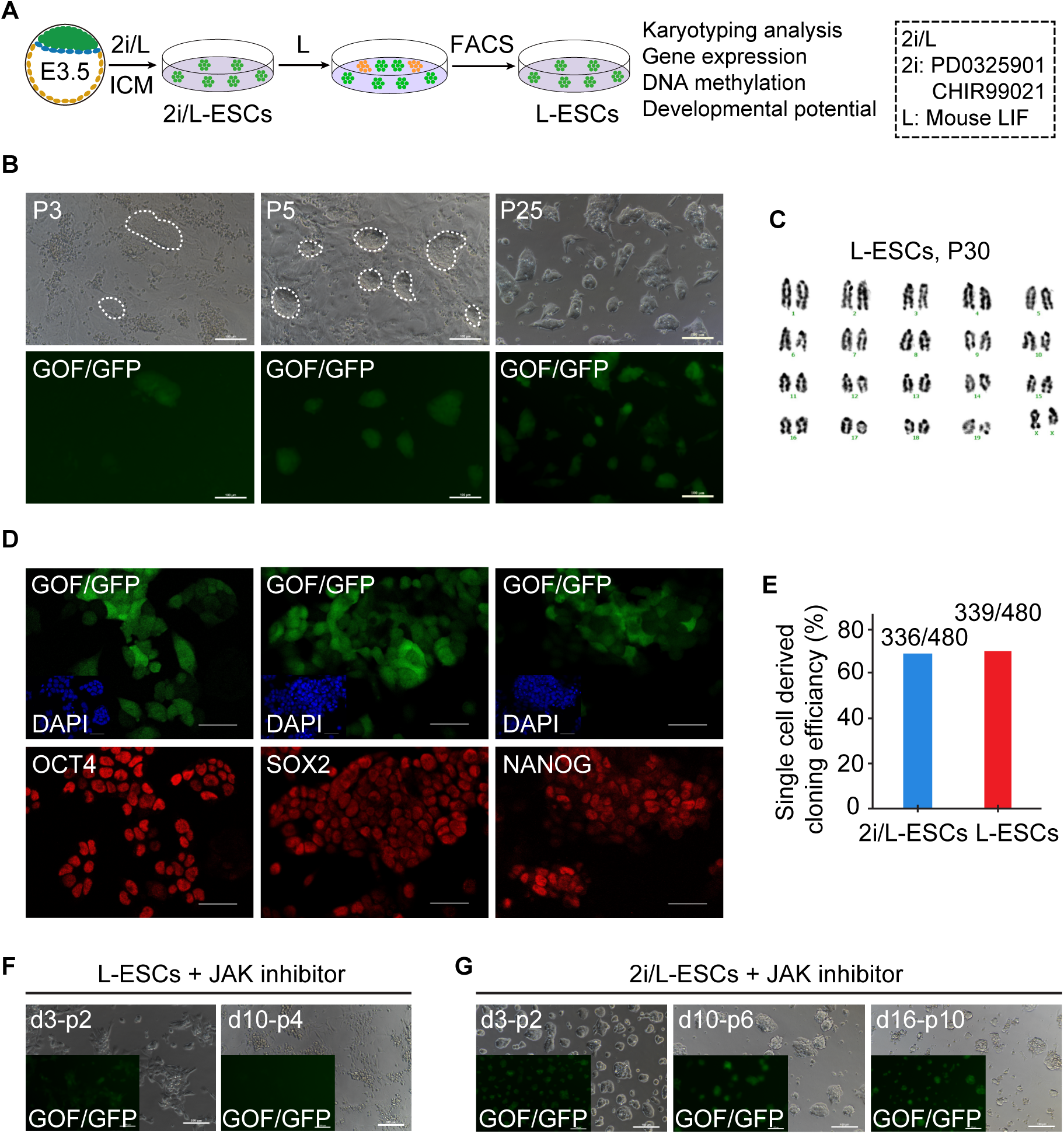
LIF alone supports ESCs self-renew and pluripotency. (A) Experimental outline of the L-ESCs derivation procedures from ESCs. (B) 2i/L-ESCs were switched to L-medium and cultured to passages 3 (p3), p5, p25. Here we use 2i/L-ESCs with GOF/GFP reporter. Scale bars, 100 μm. (C) Karyotyping of L-ESCs (p30). (D) Immunostaining of OCT4, SOX2 and NANOG in L-ESCs. Scale bars, 50 μm. (E) Single cell clonogenicity efficiency in L-ESCs and 2i/L-ESCs. (F) L-ESCs were treated with JAK inhibitor I after day 3 p2 and day 10 p4. Scale bars, 100 μm. (G) 2i/L-ESCs were treated with JAK inhibitor I after day 3 p2, day 10 p6 and p10. Scale bars, 100 μm.

Next, we performed fluorescence-activated cell sorting (FACS) on multiple L-ESCs lines and the GOF/GFP^+^ L-ESCs were cultured in L-medium. The percentage of GOF/GFP^+^ L-ESCs (passages, p14-p42) ranged from 56% to 99% in several ESCs line (*SI Appendix*, Fig. S1B). After two or more repeated FACS for each L-ESCs line (*SI Appendix*, Fig. S1B), GOF/GFP^+^ L-ESCs reached nearly 98% purity, which was similar to the control 2i/L-ESCs (*SI Appendix*, Fig. S1B). These data indicate that LIF alone can maintain FACS-purified GOF/GFP^+^ L-ESCs in undifferentiated pluripotent state (Fig. 1B and *SI Appendix*, Fig. S1B) with stable growth over 40 passages (*SI Appendix*, Fig. S1C), and high alkaline phosphatase (AP) activity (*SI Appendix*, Fig. S1D). The established L-ESCs lines have normal karyotype (Fig. 1C) and express pluripotent markers OCT4, SOX2, and NANOG, confirmed by immunofluorescence (Fig. 1D). In mouse ESCs, < 1% of cells exhibit some features of 2-cell (2C) embryos, such as the expression of 2C specific transcripts (20, 21). Interestingly, L-ESCs also retained 2C features, such as ZSCAN4 and MERVL activities demonstrated by immunostaining (Fig *SI Appendix*, Fig. S1E). It has been reported that both X chromosomes are active in female naive ESCs cells (22, 23), concurrent with this, our immunostaining showed no H3K27me3 foci in female L-ESCs, suggesting that both X chromosomes are activated (*SI Appendix*, Fig. S1F). These results suggest that L-ESCs possess most of the characteristics of 2i/L-ESCs.

For a further stringent test of the pluripotency of L-ESCs, we examined the ability of clone formation from single cell level. We observed that L-ESCs could form single cell derived colonies in chemically defined LIF alone condition with high efficiency, comparable to those from 2i/L-ESCs (Fig. 1E). Furthermore, to examine how essential LIF is in maintaining L-ESCs, we withdrew LIF and then added JAK inhibitor I, and observed significantly impaired propagation of L-ESCs with rapid differentiation (Fig. 1F). However, LIF withdrawal and JAK inhibitor addition did not affect the self-renewal of 2i/L-ESCs until passages 10 (Fig. 1G). Taken together, our results suggest that LIF is an important and essential regulator in the maintenance of L-ESCs. In contrast to the previous notion that LIF and 2i were both indispensable for ESCs self-renewal, and established unique ground state of ESCs, in this study we showed that LIF alone is capable to support ESCs for self-renewal and proliferation over passage 40.

### Global transcriptome features of L-ESCs

To examine whether L-ESCs have distinct molecular features, we carried out RNA sequencing (RNA-seq) on L-ESCs, 2i/L-ESCs, S/L-ESCs and EpiSCs. Unsupervised hierarchical clustering (UHC) and principal component analysis (PCA) showed L-ESCs close to 2i/L-ESCs (Fig. 2A and B) and appeared to be at an intermediate state between naïve ESCs and primed EpiSCs (Fig. 2A). Comparing L-ESCs and 2i/L-ESCs, L-ESCs differentially expressed genes were related to embryonic morphogenesis, cellular lipid metabolic processes, pattern specification processes, embryonic organ morphogenesis and DNA hypermethylation. Whereas 2i/L-ESCs differentially expressed genes were related to stem cell development, stem cell proliferation, WNT-protein binding, gamete generation and meiotic cell cycle phase (Fig. 2C). This shows L-ESCs display distinct molecular features for pluripotency. Interestingly, Compared with L-ESCs, 2i/L-ESCs, S/L-ESCs and EpiSCs, among differentially expressed genes (24), 3,347 genes (profile 7) were significantly high expressed in L-ESCs and 2i/L-ESCs compared with S/L-ESCs and EpiSCs (Fig. 2D). Notably, a total of 1,621 genes (profile 2) were significantly upregulated in 2i/L-ESCs, compared with L-ESCs, S/L-ESCs and EpiSCs (Fig. 2D). These RNA-seq analyses suggest that L-ESCs are in a stable state between naïve and primed pluripotency.

**Figure 2.**
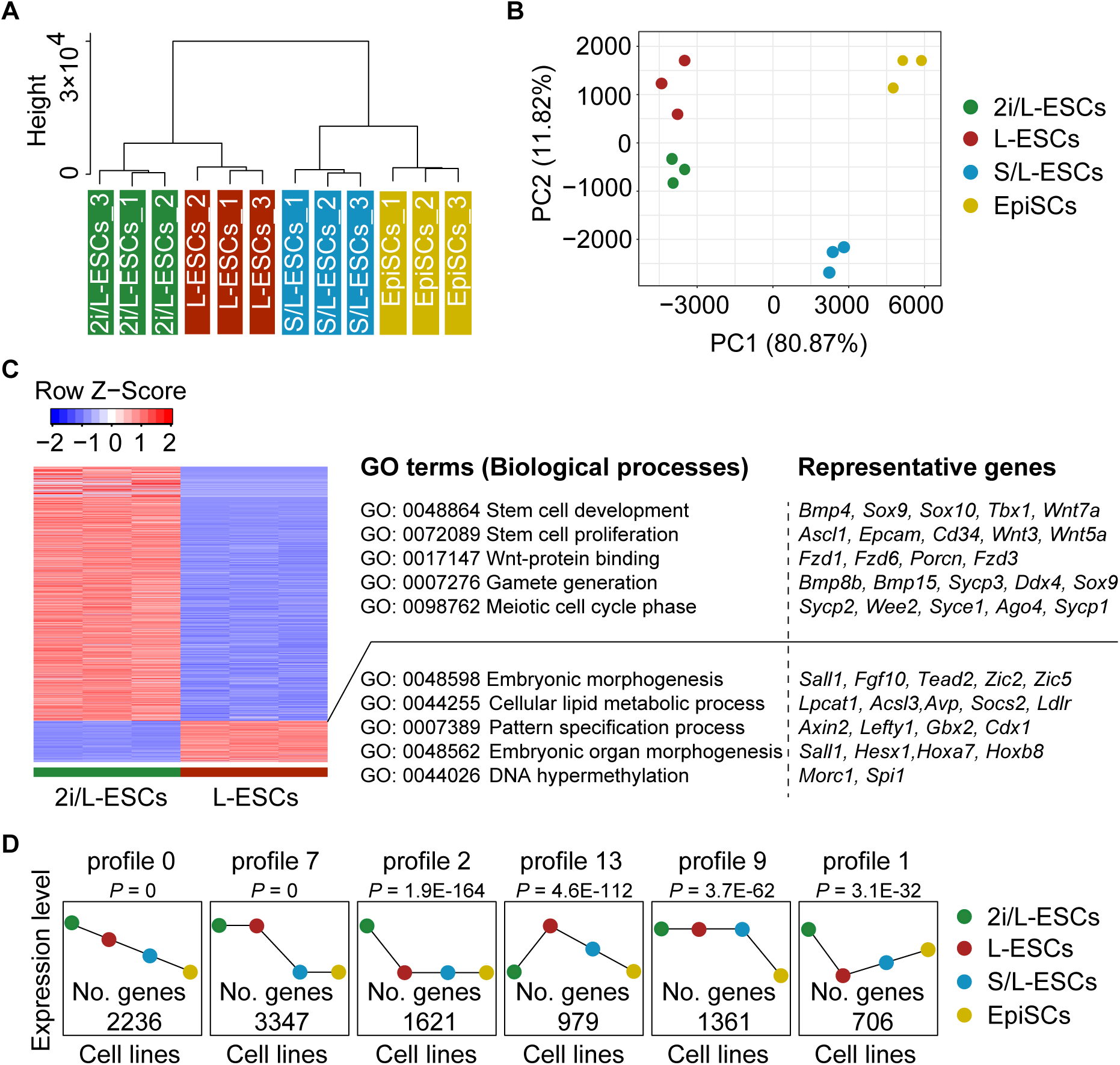
Analyses of molecular features of L-ESCs. (A) Unsupervised hierarchical clustering (UHC) of the transcriptome from three biological replicates of four pluripotent stem cell lines. (B) PCA analysis of gene expression of four pluripotent stem cells. (C) Heatmap showing differentially expressed genes (mean log2(normalized read counts) > 2, log2(fold change) > 2, adjusted *p*-value < 0.05) in L-ESCs compared with 2i/L-ESCs. Significantly enriched GO terms and representative genes in each cluster are listed on the right. (D) Compared with L-ESCs, 2i/L-ESCs, S/L-ESCs and EpiSCs, among differentially expressed genes, a total of 3,347 genes (profile 7) were significantly high expressed in L-ESCs and 2i/L-ESCs compared with S/L-ESCs and EpiSCs; a total of 1,621 genes (profile 2) were significantly upregulated in 2i/L-ESCs, compared with L-ESCs, S/L-ESCs and EpiSCs.

### L-ESCs exhibit DNA hypermethylation and reserve genomic imprints

ESCs cultured in 2i/LIF or in LIF plus serum supplemented media represent two states of pluripotency of ESCs. Despite their similarities in pluripotency, 2i/L and S/L-ESCs rely on different signaling pathways and display strong differences in transcriptional and epigenetic landscapes (25–27). Here, we asked whether there are different epigenetic marks among L-ESCs, 2i/L-ESCs and S/L-ESCs. Whole-genome bisulfite sequencing (WGBS) was performed and DNA methylation profiling of L-ESCs with 2i/L-ESCs and S/L-ESCs was compared. The levels of DNA methylation in L-ESCs (median CpG methylation of ∼80%) were comparable to S/L-ESCs (median ∼90%) and higher than 2i/L-ESCs (median ∼30%) (Fig. 3A). This DNA methylation occurs across most methylated regions including intragenic, intergenic, exon, intron, short and long interspersed nuclear elements (SINEs and LINEs, respectively) and long terminal repeats (LTRs) (*SI Appendix*, Fig. S2A). Additionally, expression of DNA methylation associated genes was assessed using qPCR. As expected, DNA methyltransferases *Dnmt3a* and *Dnmt3l* was significantly upregulated in GOF/GFP positive L-ESCs compared with GOF/GFP negative cells from the L-ESCs reprogramming process (*SI Appendix*, Fig. S2B). Moreover, the transcriptional level of genes known to influence DNA methylation levels, such as *Prdm14* and *Nanog* were significantly downregulated in L-ESCs (*SI Appendix*, Fig. S2B).

**Figure 3.**
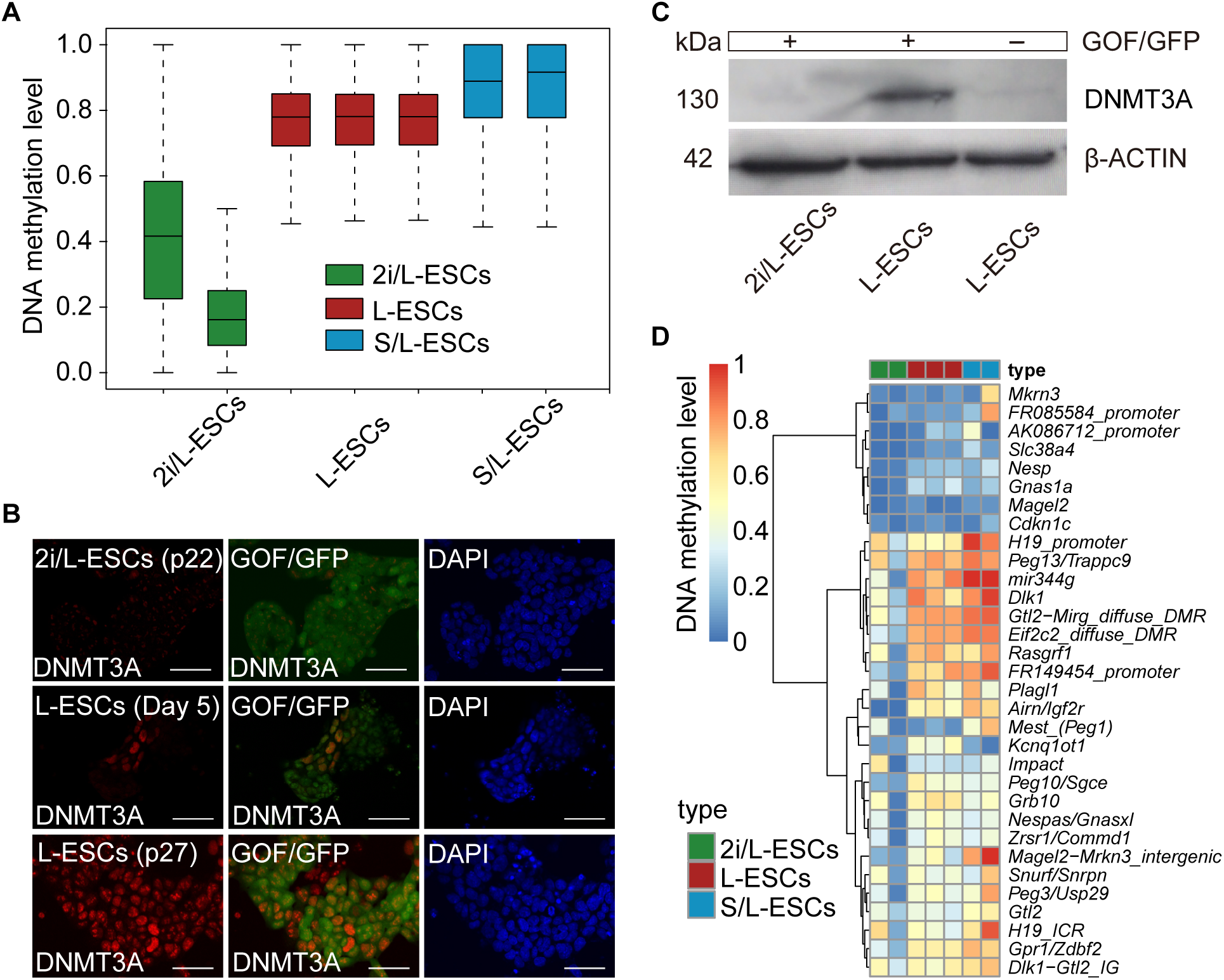
DNA methylation pattern of L-ESCs. (A) DNA methylation level of 2 kilobase (kb) genomic tiles. Source data are provided in Table S1. (B) Immunostaining of Dnmt3a in 2i/L-ESCs and different passages L-ESCs. Scale bars, 50 μm. (C) Western blotting analysis for Dnmt3a in early reprogramming stage (day 5) L-ESCs (GOF/GFP positive and negative cells). (D) Heatmap showing DNA methylation level of ICRs in three different stem cells.

Next, we examined the dynamic changes of DNMT3A level in the process of L-ESCs reprogramming. Interestingly, the protein level of DNMT3A was high in early reprogramming stage (day 5) GOF/GFP positive L-ESCs (Fig. 3B and C). Upon withdrawal of PD0325901 and CHIR99021, heterogeneous expression of DNMT3A was detected in nuclei of L-ESCs reprogramming at day 5, and in long-term culture the DNMT3A protein level was significantly increased in p27 stage L-ESCs (Fig. 3B), consistent with the higher methylation in L-ESCs. These data is also consistent with the notion that PD0325901 promotes downregulation of DNA methylation (15, 16). The results showed DNMT3A is important factor to regulate DNA methylation in L-ESCs which possess hypermethylation state.

Proper genomic imprinting is essential for embryonic development (28, 29). We further performed genomic imprinting analysis on L-ESCs, S/L-ESCs and 2i/L-ESCs. Notably, compared with 2i/L-ESCs, the DNA methylation levels at imprinting control regions (ICRs) were markedly higher in L-ESCs and were similar to S/L-ESCs (Fig. 3D). Collectively, we conclude that L-ESCs exhibited global genomic hypermethylation and reserved genomic methylation in the majority of imprinting control regions.

### Serum treatment increase the efficiency of LIF-dependent ESCs reprogramming

Since the S/L-ESCs possess high levels of DNA methylation (25), we next asked if serum treatment (prior to reprogramming of L-ESCs) may enhance the DNA methylation and then increase the efficiency of LIF dependent L-ESCs reprogramming. We switched 2i/L-ESCs to S/L-medium for five days of induction, then S/L-ESCs were cultured in LIF only (L-medium) to assess the LIF dependent L-ESCs reprogramming. Our result indicates that S/L induction for 5 days significantly increased the number of AP^+^ colonies compared with 2i/L-ESCs (Fig. 4A and *SI Appendix*, Fig. S3A). Consistent with this, 1×10^5^ cells were seeded into 24-well cell culture plate in L-medium, the number of GOF/GFP^+^ colonies obtained from S/L induction group compared with 2i/L-ESCs was drastically increased (Fig. 4B). To confirm this, flow cytometry analysis showed that the percentage of GOF/GFP^+^ cells in the S/L induction group was also increased compared with 2i/L-ESCs (Fig. 4C). Furthermore, we tested this reprogramming process of ASCs in LIF alone medium using our previously published hypermethylated ASCs (17, 30) and showed that ASCs can also be efficiently reprogrammed into LIF-dependent ESCs using L-medium (*SI Appendix*, Fig. S3B and C).

**Figure 4.**
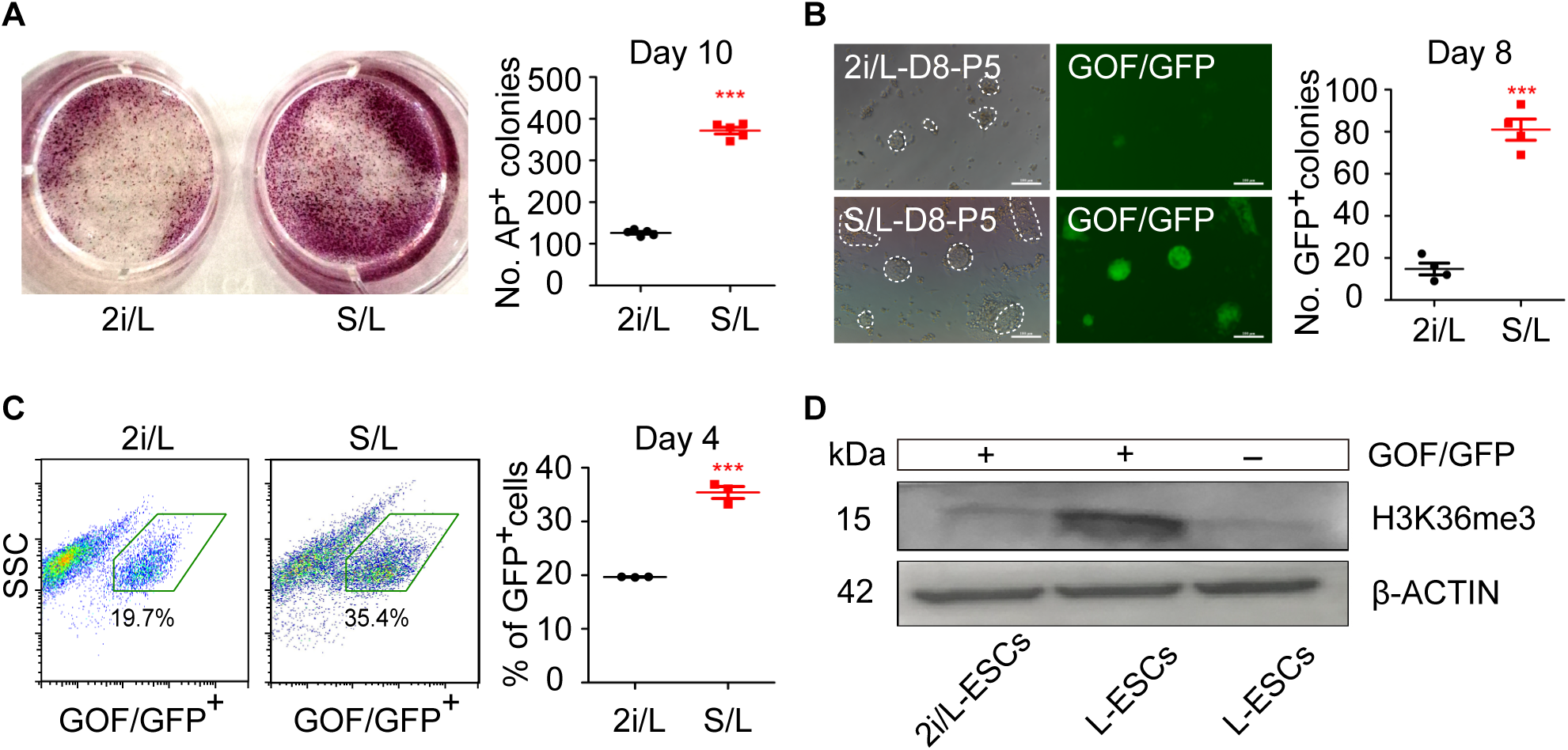
Serum improves the efficiency of L-ESCs reprogramming. (A) Left: Alkaline phosphatase (AP) staining on 2i/L-ESCs and S/L-ESCs (2i/L-ESCs were cultured in S/L medium for 5 days) were switched to L-medium and after 10 days culture. Right: Quantification of number of AP positive colonies after 10 days culture. Error bars are mean ± SD (n = 5). *P* values were calculated by two tailed Student’s *t*-test, *p* < 0.05. (B) Left: GOF/GFP positive colonies on 2i/L-ESCs and S/L-ESCs (2i/L-ESCs were cultured in S/L medium for 5 days) were switched to L-medium and after 8 days culture. Scale bars, 100 μm. Right: Quantification of number of GOF/GFP positive colonies after 8 days culture. Error bars are mean ± SD (n = 4). *P* values were calculated by two tailed Student’s *t*-test, *p* < 0.05. (C) Left: Fluorescence-activated cell sorting (FACS) based on GOF/GFP positive cells, after 2i/L-ESCs and S/L-ESCs (2i/L-ESCs were cultured in S/L medium for 5 days) were switched to L-medium and after 4 days culture. Right: Quantification of Percentage of GOF/GFP positive cells after 4 days culture. Error bars are mean ± SD (n = 3). *P* values were calculated by two tailed Student’s *t*-test, *p* < 0.05. (D) Western blotting analysis for H3K36me3 in early reprogramming stage (day 5) L-ESCs (GOF/GFP positive and negative cells).

### DNA methylation is indispensable for L-ESCs self-renewal

Next, we asked whether DNA methylation is critical for this reprogramming, and investigated roles of different DNA methyltransferase in the early reprogramming processes. We separated GOF/GFP^+^ and GOF/GFP^−^ L-ESCs from early reprogramming processes by FACS. As expected, *Dnmt3a* and *Dnmt3l* expression levels in GOF/GFP^+^ L-ESCs were significantly higher than in GOF/GFP^−^ L-ESCs (*SI Appendix*, Fig. S2B), as well as DNMT3A protein level (Fig. 3C). Interestingly, we also found higher expression level of H3K36me3 in GOF/GFP^+^ early reprogramming stage (day 5) L-ESCs (Fig. 4D). This result is consistent with recent reports of H3K36me3 as a guard for the DNA methylation process (31).

To unequivocally demonstrate whether stable L-ESCs self-renewal depends on DNA methylation, we next examined the role of DNA methyltransferases (DNMTs) on the regulating L-ESCs self-renewal processes by the DNMT inhibitor 5-aza-2’-deoxycytidine (5-Aza). 5-Aza has been widely used as a DNA methylation inhibitor to experimentally induce gene expression and cellular differentiation (32, 33). We cultured 2i/L-ESCs and L-ESCs in their respective medium with 5-Aza and observed morphological changes of both 2i/L-ESCs and L-ESCS. 5-Aza treated 2i/L-ESCs retained typical dome-shaped clonal morphology and were able to stably propagate at least ten passages (Fig. 5A and *SI Appendix*, Fig. S4A). In addition, there were slight changes in the expression level of pluripotent genes (including *Nanog, Sox2* and *Prdm14*) between 5-Aza treated 2i/L-ESCs and untreated 2i/L-ESCs (*SI Appendix*, Fig. S3B). However, 5-Aza treated L-ESCs failed to maintain its self-renewal. There were few GOF/GFP^+^ L-ESCs which survived after seven days upon 5-Aza treatment and cells underwent apoptosis eventually (Fig. 5B). These data indicate that L-ESCs are differentially sensitive to inhibition of DNA methyltransferase by 5-Aza compared with 2i/L-ESCs.

**Figure 5.**
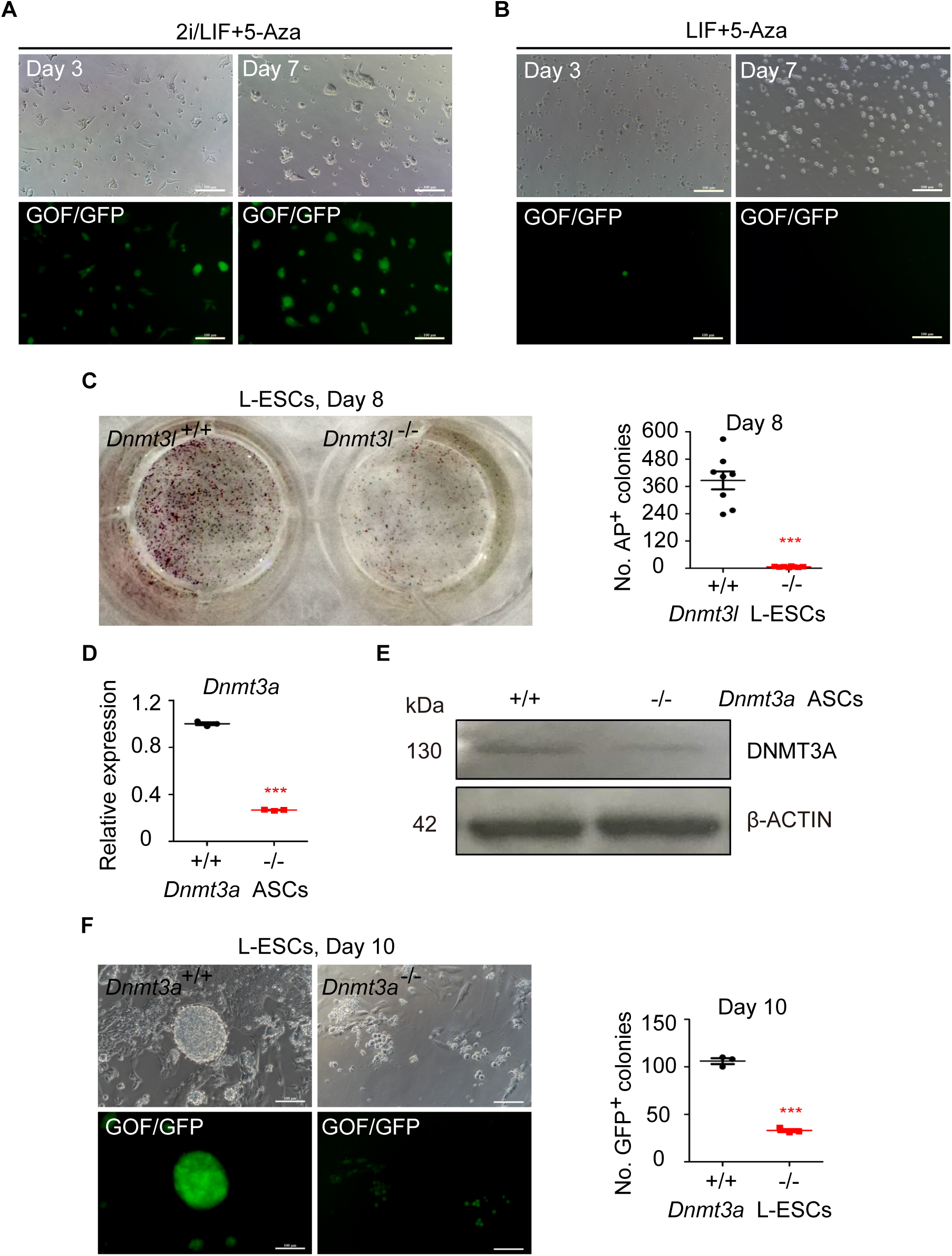
DNA methylation is indispensable for L-ESCs self-renew. (A) 2i/L-ESCs were treated with 5-Aza after 3 and 7 days, 2i/L-ESCs retained typical dome-shaped clonal morphology. Scale bars, 100 μm. (B) L-ESCs were treated with 5-Aza after 3 and 7 days, there was a few GOF/GFP^+^ L-ESCs survived after 7 days 5-Aza treatment and to apoptosis in final. Scale bars, 100 μm. (C) Left: AP staining on wild type ESCs and *Dnmt3l*^−/-^ ESCs were switched to L-medium and after 8 days culture. Right: Quantification of number of AP positive colonies after 8 days culture. Error bars are mean ± SD (n = 8). *P* values were calculated by two tailed Student’s *t*-test, *p* < 0.05. (D) Relative expression of *Dnmt3a* by qPCR in *Dnmt3a^−/-^* ASCs and *Dnmt3a^+/+^* ASCs. Error bars are mean ± SD (n = 3). *P* values were calculated by two tailed Student’s *t*-test, *p* < 0.05. (E) Western blotting analysis for DNMT3A in *Dnmt3a^−/-^*-ASCs and *Dnmt3a^+/+^*-ASCs. (F) Left: GOP/GFP positive colonies on wild type ASCs and *Dnmt3a*^−/-^ ASCs were switched to L-medium and after 10 days culture. Right: Quantification of number of GOP/GFP positive colonies after 10 days culture. Error bars are mean ± SD (n = 8). *P* values were calculated by two tailed Student’s *t*-test, *p* < 0.05.

To further investigate the important role of DNA methylation on LIF-dependent ESCs reprogramming processes, we used *Dnmt3l* knockout ESCs (*Dnmt3l^−/-^*-ESCs) which were cultured in S/L medium (Fig. 5C) and generated *Dnmt3a* knockout ASCs line (*Dnmt3a^−/-^*-ASCs) which were cultured in ABC/L medium (Fig. 5D and E) (17) and then switched to chemically defined LIF medium. As expected, both *Dnmt3l* and *Dnmt3a* knockout cells significantly reduced the efficiency of LIF-dependent ESCs reprogramming (Fig. 5C and F). Whereas wild type ESCs and ASCs derived L-ESCs displayed normal self-renew and proliferation, the proliferation of *Dnmt3l^−/-^* and *Dnmt3a^−/-^* L-ESCs decreased dramatically (Fig. 5C and F; *SI Appendix*, Fig. S4A and C). Taken together, our data demonstrate that DNA hypermethylation promotes the induction of LIF-dependent ESCs reprogramming.

### *In vitro* and *in vivo* differentiation ability of L-ESCs

An important criterion for pluripotent ESCs is the ability to differentiate *in vitro* and *in vivo* (34). Upon 2i and LIF withdrawal, pluripotent ESCs differentiate into three germ layers, mesoderm, endoderm, and ectoderm (35). We cultured 2i/L-ESCs and L-ESCs in N2B27 basic medium only, without 2i/L and LIF. In these culture conditions, the ESCs differentiated. After 3-day and 6-day differentiation, we performed quantitative qPCR analysis and immunostaining. Interestingly, after 3-day differentiation, the expression level of all mesoderm, endoderm, and ectoderm genes were significantly increased in L-ESCs compared with 2i/L-ESCs (Fig. 6A). Compared with 3-day differentiation, 6-day culturing significantly increased mesoderm, endoderm, and ectoderm genes expression level in 2i/L-ESCs but not in L-ESCs (*SI Appendix*, Fig. S5A and B). This indicates that L-ESCs have strong flexibility and differentiation ability depends on the environment changes. Nevertheless, 6-day differentiation ability between 2i/L-ESCs and L-ESCs was not significantly different (*SI Appendix*, Fig. S5C). Furthermore, we confirm protein levels of mesoderm, endoderm and ectoderm markers by immunostaining (Fig. 6B). In addition, similar to 2i/L-ESCs, L-ESCs also generated teratomas that contained derivatives of the three germ layers (Fig. 6C). The results showed that L-ESCs have differentiation ability both *in vitro* and *in vivo*, and is able to express important differentiation genes in a shorter space of time when compared to 2i/L-ESCs.

**Figure 6.**
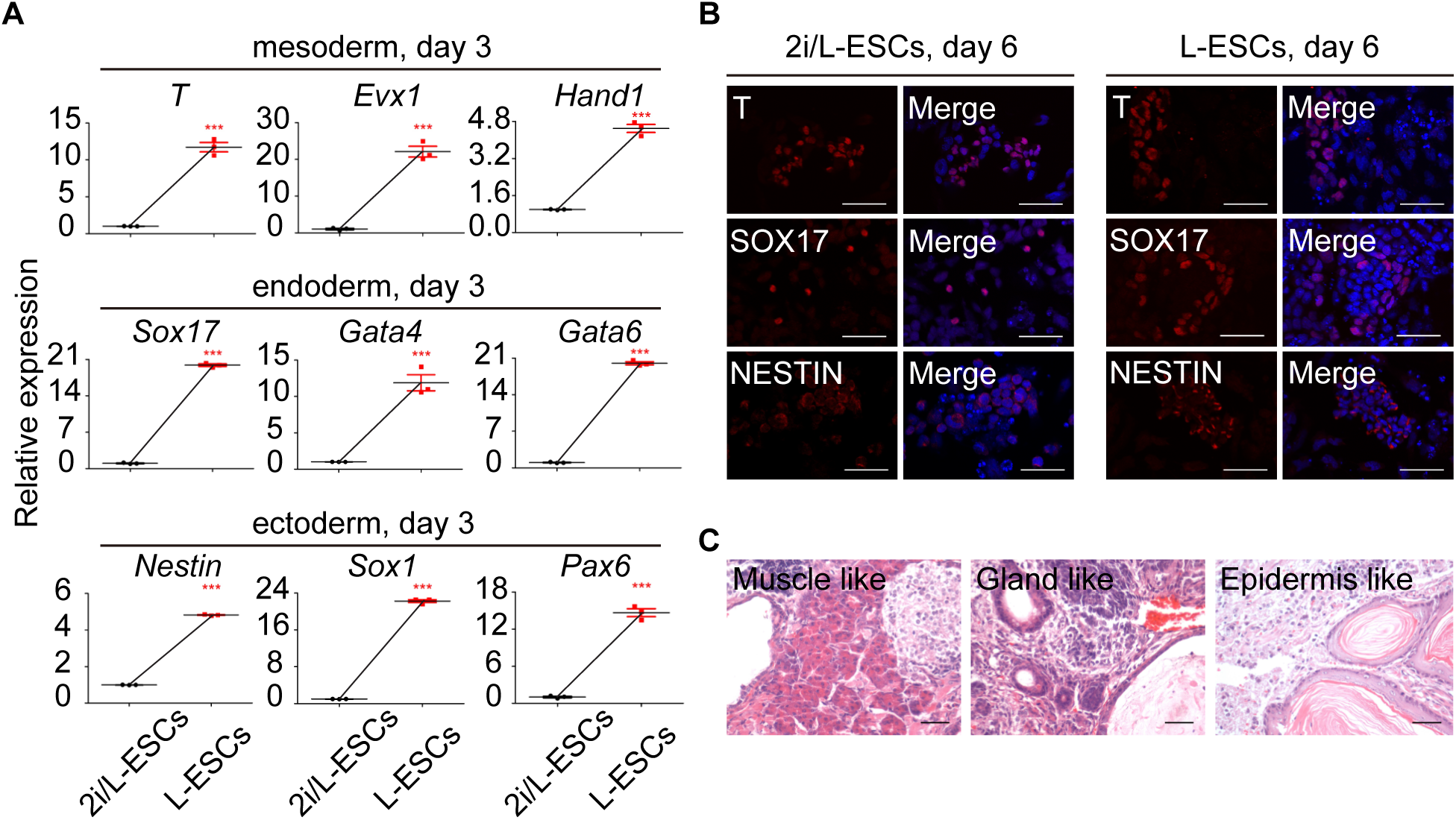
The pluripotency of L-ESCs *in vivo* and *in vitro*. (A) Relative expression of mesoderm, endoderm and ectoderm genes measured by qPCR, after L-ESCs were 3 days *in vitro* differentiation. Error bars are mean ± SD (n = 3). *P* values were calculated by two tailed Student’s *t*-test, *p* < 0.05. (B) Immunostaining of T, SOX17 and NESTIN, after 2i/L-ESCs and L-ESCs were 6 days *in vitro* differentiation. Scale bars, 50 μm. (C) Mature teratomas from L-ESCs. Left: mesoderm, muscle like cells; middle: endoderm, gland like cells; right: ectoderm, epidermis like cells. The sections were stained with haematoxylineosin. Scale bars, 50μm.

### Contribution of L-ESCs to full-term embryonic development

Finally, we tested the *in vivo* developmental potential of L-ESCs in chimeric embryos. Using L-ESCs derived from 2i/L-ESCs, we injected L-ESCs into 8-cell stage embryos (Fig. 7A). We noticed that L-ESCs successfully integrated into E13.5 germlines of chimeras. Notably, 36.8% (7/19) of recovered embryos showed chimeric contribution and 57.1% (4/7) of chimeric embryos displayed germlines contribution (Fig. 7B and C). We further tested whether it is possible to obtain L-ESCs-derived postnatal chimeric mice. Of all 20 born pups, 5 L-ESCs-derived chimeras (25%) were obtained (Fig. 7D and E). Hence, these data demonstrate the pluripotency of L-ESCs and their chimeric competency to both germlines contribution and full-term development.

**Figure 7.**
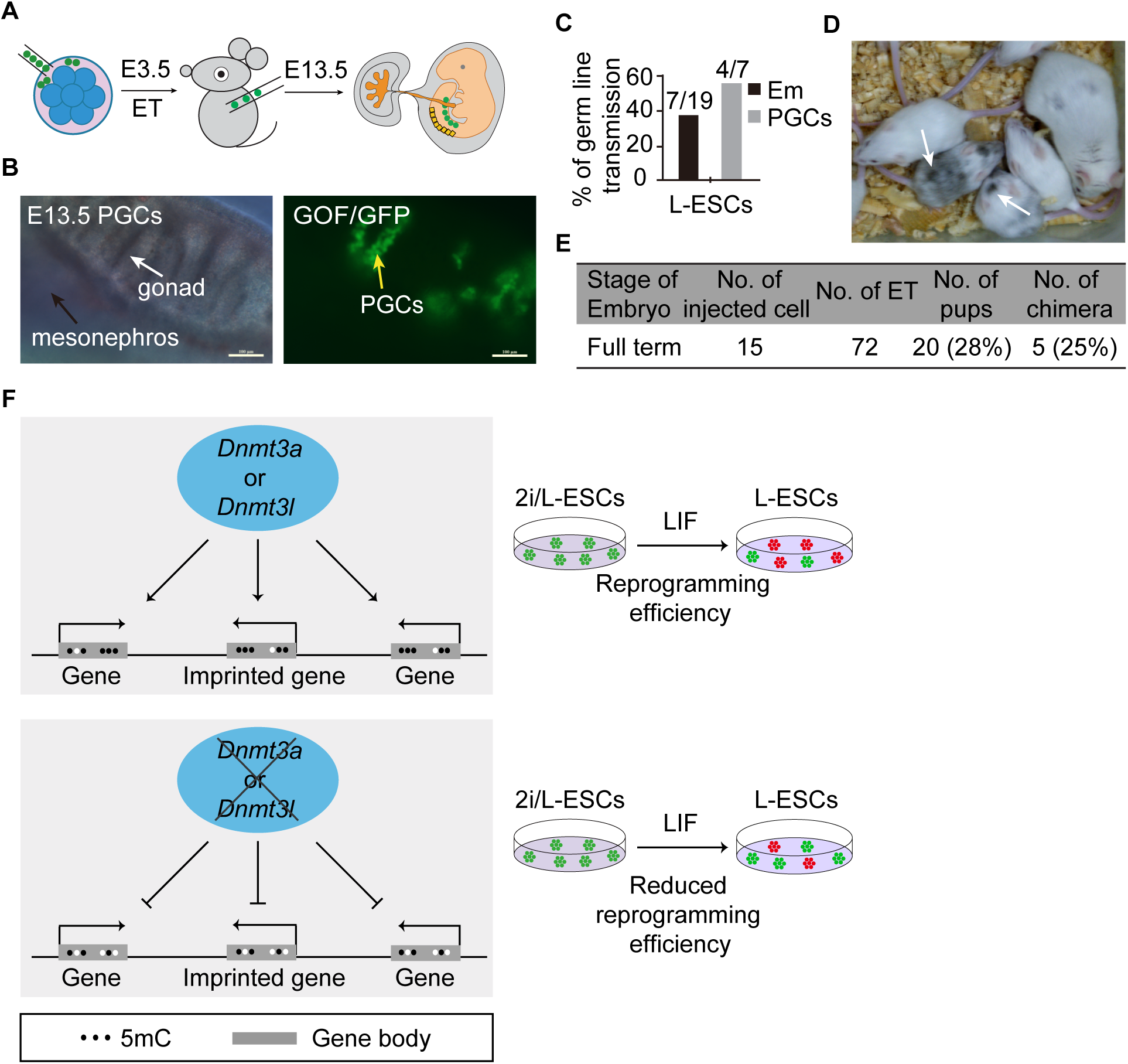
Ability of L-ESCs to full-term embryonic development. (A) Schematic of eight cell embryos injection protocol. (B) Germline transmission of L-ESCs in E13.5 chimeras. PGCs were shown by GOF/GFP-positive cells (arrow). black arrow: mesonephros; white arrow: gonad; yellow arrow: gonadal PGCs. Scale bars, 100 μm. (C) Summary of E13.5 chimera assays by L-ESCs injection. The black bar chart shows the percentages of chimeras among the collected E13.5 conceptuses, embryonic tissues (Em); gray bar, integration into primordial germ cells (PGCs) among the recovered E13.5 chimeras. (D) Chimeric pups generated by injecting L-ESCs in ICR host blastocysts. (E) The summary of full term chimeric pups were derived by L-ESCs. (F) Schematic of DNA methylation affects LIF dependent embryonic stem cells reprogramming process.

## Discussion

ESCs are derived from the inner cell mass (ICM) of the blastocyst, and self-renew indefinitely *in vitro* (4, 6). The signaling of WNT, ERK and JAK/STAT3 are main regulators that combine to control pluripotency, however, precise function of the individual signaling pathways is unclear (7). In this study, we represent the induction of one novel cell type, L-ESCs from 2i/L-ESCs, which depend on JAK/STAT3 signaling alone, and provide new insights on the nature of pluripotent stem cells. In particular, the L-ESCs show higher DNA methylation levels than 2i/L-ESCs (Fig. 3A), and based on transcriptional level, L-ESCs appeared to be at an intermediate state between naïve ESCs and primed EpiSCs (Fig. 2A). We also find that genomic imprints are more stable in L-ESCs relative to 2i/L-ESCs (Fig. 3D). Based on the gene expression and DNA methylome analysis, L-ESCs appeared to be at an intermediate state between naïve ESCs and primed EpiSCs, and may represent stable cells with the characteristics of the early postimplantation epiblast.

LIF signaling include JAK/STAT, MARK and PI(3)K pathways, and stimulates a states of self-renewal, and determines the fate of cells (7). In mouse ESCs, it is generally believed that LIF signaling is skewed towards survival and self-renewal, whereas activation of canonical WNT signaling and blockade of FGF/ERK blocks cell differentiation (7). In this study we show that under LIF alone medium, some proportion of surviving ESCs acquires new features. These L-ESCs maintained self-renewal and pluripotency over passage 40. We show that L-ESCs died in 10 days in medium with JAK inhibitor (Fig. 1F and G). It has been clear that LIF is critical to L-ESCs self-renewal and to maintain undifferentiated state. One hypothesis is that 2i/L-ESCs cultured in L-medium became heterogeneous, majority of 2i/L-ESCs differentiation in this regime, and only small proportion indicates the presence of naïve ESCs, which the JAK/STAT3 may favor to bind to cofactors or intrinsic factor that promote self-renew. Recently Ying et al reported STAT3 signaling functions in a binary “on/off” manner, however they used S/L medium, the defined mechanism needs to be further explored (36).

DNA methylation is of paramount importance for mammalian embryonic development and DNA methylation deficient embryos die at such an early stage of development (37). Here, we show that DNA hypermethylation increased the efficiency of L-ESCs reprogramming in S/L medium, whereas *Dnmt3a* and *Dnmt3l* knockout model and 5-Aza treatment affect the efficiency of inducing L-ESCs reprogramming and self-renewal. Interestingly, triple-knockout (TKO) mouse S/L-ESCs for *Dnmt1, Dnmt3a* and *Dnmt3b* exhibit DNA hypomethylation, grows robustly and maintains their undifferentiated characteristics (38). Unlike mouse ESCs, conventional „primed‟ human ESCs cannot tolerate *Dnmt1* deletion, emphasizing the functional differences between mouse and human ESCs (39). We suggest that embryonic stem cells cultured in LIF alone exhibit media dependent DNA hypermethylation and this state support L-ESCs self-renew and proliferation. Notably, LIF-dependent ESCs reprogramming efficiency is significantly reduced in *Dnmt3a* or *Dnmt3l* knockout ESCs (Fig. 7F). We also show that DNMT3A and H3K36me3 expression were higher in L-ESCs compare to 2i/L-ESCs. Recently, multiple studies suggested that H3K36me3 participates in cross-talk with other chromatin marks, and promotes de novo DNA methylation by interacting with DNMTs and SETD2 (31). H3K36me3 is responsible for establishing and safeguarding the maternal epigenome (31). Our result showed that H3K36me3 and DNMT3A were highly expressed in L-ESCs, supports this hypothesis.

Epigenetics including genomic imprinting has widespread roles in mammals, affecting embryonic and placental development and transmission of nutrients to the fetus, and regulating critical aspects of mammalian physiology, such as metabolism, neuronal development and adult behavior (28). We show that L-ESCs reserve hypermethylated imprinting genes, easily differentiate in medium without LIF, which may suggest unique features for ESCs pluripotency. On the other hand, unlike ASCs with high development potency in chimeras, a single L-ESCs do not contribute to development of the embryo to such an extent, suggesting that L-ESCs state is an intermediate between naïve ESCs and primed EpiSCs, and its pluripotency are more close to S/L-ESCs and EpiSCs. In conclusion, this study demonstrates LIF alone is capable to support mouse ESCs reprogramming and provides mechanistic insight into the role of global DNA (de)methylation.

## Materials and Methods

### Ethics statement

Animal care and use were conducted in accordance with the guidelines of Inner Mongolia University, China. Mice were housed in a temperature-controlled room with proper darkness-light cycles, fed with a regular diet, and maintained under the care of the Laboratory Animal Unit, Inner Mongolia University, China. The mice were sacrificed by cervical dislocation. This study was specifically approved by the Institutional Animal Care and Use Committee, Inner Mongolia University, China. Oct4-**Δ**PE-GFP (GOF/GFP) transgenic mice (19) were used here with a mixed background of MF1, 129/sv, and C57BL/6J strains.

### Derivation of 2i/L-ESCs

Mouse embryos blastocysts (E3.5) were isolated from 129/sv females mated with GOF/GFP transgenic males. Green fluorescence indicated that GFP expression of the reporter is under the control of *Oct4* promoter and distal enhancer. This GFP transgene shows expression in the ICM of blastocysts and PGC *in vivo*, and in ESCs (19). ESCs culture medium consists of N2B27 medium (Life technology) supplemented with PD0325901 (PD, 1 μM, Miltenyi Biotec), CHIR99021 (CH, 3 μM, Miltenyi Biotec) and leukemia inhibitory factor (LIF, 1000 IU/ml, Millipore), henceforth were called 2i/L medium. Zona pellucida of blastocysts were removed by Acidic Tyrode‟s Solution (Sigma-Aldrich), and then placed to 24-well fibronectin-coated (FN, 16.7 μg/ml, Millipore) plate with 2i/L medium. ICM of blastocysts cultures grew efficiently and formed outgrowing colonies in 5-7 days culture. The resulting colonies were further cutting into smaller pieces by glass needles after 5-7 days culture, and then the colonies passaged by Accutase (Life technology) regularly on at every 2 days interval.

### Derivation of L-ESCs

1×10^5^ 2i/L-ESCs were switched on fibronectin-coated (16.7 μg/ml, Millipore) 24-well cell culture plate containing L-medium which are N2B27 medium supplemented with leukemia inhibitory factor (1000 IU/ml, Millipore), and we call these cells as L-ESCs. Dependent on cell growth, L-ESCs were passage every other day in the early stage. After being cultured for 4-5 passages or 14-42 passages, GOF/GFP positive and negative L-ESCs were purified by flow-cytometry sorting by BD FACSAria (BD Biosciences) and further analysis. GOF/GFP positive purified L-ESCs were passage every other day treated with Accutase (Life technology). L-ESCs were capable of self-renewal for over 40 passages. For inhibitor treatment experiment, we added JAK inhibitor I (0.6 μM, Calbiochem) or 5-Aza (6 μM, Sigma) into L-ESCs culture medium.

### Derivation of S/L-ESCs

2i/L-ESCs were switch to fibronectin-coated plate with standard ES medium (Knockout DMEM; Knockout Dulbecco‟s modified Eagle‟s medium) supplemented with 20% fetal calf serum, 0.1 mM 2-mercaptoethanol, 2 mM L-glutamine, 0.1 mM non-essential amino acid, 50 U/ml Penicillin/Streptomycin and 1000U/ml LIF without feeder cells, we named these cells as S/L-ESCs.

### Cell differentiation

2i/L-ESCs and L-ESCs were cultured in N2B27 medium for 3 to 6 days withdrawal of PD0325901, CHIR99021 and LIF, and LIF respectively.

### Colony formation assay

Single 2i/L-ESCs and L-ESCs were seeded at a fibronectin-coated 96-well plates using mouth pipette, containing 2i/L and L-medium, respectively. The cells were cultured for 10 days and the number of colonies was assessed.

### Western blot

Cells were collected with Accutase (Life technology), washed three times with DPBS, and lysed in buffer that contained 20 mM Tris (pH 8.0), 137 mM NaCl, 100 g/l glycerol, 50 g/l Triton X-100, and 4 g/l EDTA; 1 μl PMSF (0.1 M) and 10 μl phosphatase inhibitor (10 g/l) were added per 1 ml lysis buffer immediately before use. Proteins were denatured with 2 × SDS at 95 °C for 5 min. A total of 20 μg denatured protein was run on 8% or 10% SDS– PAGE gel and transferred to polyvinylidene difluoride (PVDF) membrane. Membranes were blocked with 5% nonfat milk in 1 × TBS with 0.05% Tween-20 (TBST) for 1h. Samples were probed with primary antibodies overnight at 4°C. The primary antibodies used were anti-DNMT3A (CST, 3598S; dilution 1:1,000), anti-H3K36me3 (Abcam, ab9050; working concentration, 1 μg/ml), and anti-β-ACTIN (Abcam, ab8227; dilution 1:5,000). Blots were rinsed with TBST. Membranes were incubated with HRP-conjugated secondary antibodies for 60 min at room temperature, and proteins were detected by ECL plus reagent. After rinsing with TBST, Clarity^TM^ Western ECL Substrate (BIO-RAD) was used for visualization, and ChemiDoc^TM^ MP Imaging System (BIO-RAD) was used for band detection.

### Alkaline phosphatase (AP) staining

AP staining was carried out using AP staining kit from Sigma (86R-1KT) according to manufacturer‟s instructions. Briefly, the cells were fixed by 4% paraformaldehyde for 10 min, and then were stained by AP staining solution for overnight at room temperature.

### Immunostaining

Cultured ESCs were briefly washed with PBS and fixed in 4% paraformaldehyde in PBS for 15 min at room temperature. Cells were permeabilized for 30 min with 1% BSA and 0.1% Triton X-100 in PBS. Antibody staining was carried out in the same buffer at 4°C for overnight. The slides were subsequently washed three times in 1% BSA, 0.1% Triton X-100 in PBS (5 min each wash), were incubated with secondary antibody for 1h at room temperature in the dark, washed once for 5 min in 1% BSA, 0.1% Triton X-100 in PBS and twice for 5 min in PBS. The slides were then mounted in Vectashield with DAPI (Vector Laboratories) and imaged using a Olympus FV1000 confocal microscope. Primary antibodies used were: anti-OCT4 (BD Biosciences, Catalog Number: 611203, 1:200), anti-NANOG (eBioscience, Catalog Number: 14-5761, 1:500), anti-SOX2 (Santa cruz, Catalog Number: sc-17320, 1:200), anti-H3K27me3 (Upstate, Catalog Number: 07-449, 1:500), anti-ZSCAN4 (Abcam, Catalog Number: ab106646, 1:200), anti-MERVL (HuaAn Bio, Catalog Number: ER50102, 1:100), anti-DNMT3A (abcam, Catalog Number: ab79822, 1:500), anti-NESTIN (BOSTER Bio, Catalog Number: BM4494, 1:50), anti-BRACHYURY (R & D Systems, Catalog Number: AF2085, 1:100) and anti-SOX17 (R & D Systems, Catalog Number: AF1924, 1:100). All secondary antibodies used were Alexa Fluor highly crossed adsorbed (Molecular Probes).

### Flow cytometry

GOF/GFP ESCs were harvested by Accutaes and sorting by BD LSRFortessa. Green fluorescence indicated that GFP expression of the reporter is under the control of Oct4 promoter and distal enhancer. This GFP transgene shows expression in the ICM of blastocysts and PGCs in vivo, and in ESCs. No GOF/GFP ESCs were used for FACS gating negative control. Measure fluorescence (detector 488 nm channel for GFP) by flow cytometer. Gating out of residual cell debris and measure diploid and tetraploid DNA peaks. A region representing GFP-positive cells were used to identifiy living cells and collected.

### Teratomas formation

The 2i/L-ESCs and L-ESCs were disaggregated using Accutase, and 1×10^6^ cells were injected into under epithelium of NOD–SCID mice. Three to five weeks after transplantation, tumor(s) were collected and fixed with 4% paraformaldehyde, and processed for paraffin sectioning. Sections were observed following Hematoxylinand Eosin staining.

### Karyotyping

ESCs were prepared for cytogenetic analysis by treatment with colcemid (Sigma) at a final concentration of 0.1 μg/ml for 3h to accumulate cells in metaphase. Cells were then exposed to 0.075 M KCl for 25 min at 37°C and fixed with 3:1 methanol: acetic acid. Air-dried slides were generated and G-banded following standard GTG banding protocols.

### Production of chimeras

8-10 ESCs were injected gently into the ICR mice eight-cell stage embryos using a piezo-assisted micromanipulator attached to an inverted microscope. The injected embryos were cultured in KSOM medium (Millipore) at 37°C in a 5% CO_2_ atmosphere for overnight and then transferred to the uteri of pseudopregnant ICR mice at 2.5 days post coitus (dpc). The embryos were isolated at embryonic stage E13.5 and check germline transmission. Full term chimeras were confirmed by the coat color pattern of the pups at birth.

### Real-Time PCR

Total RNA was isolated with the RNeasy Plus Mini Kit (Qiagen) and reverse transcribed into cDNA using the Reverse Transcription System (Promega) according to the manufacturer‟s instructions. Quantitative real-time PCR (qRT-PCR) was conducted using a LightCycler® 96 Instrument (Roche Molecular Systems) and qRT-PCR reaction was performed with KAPA SYBR FAST qPCR kit (KAPA Biosystems). At least triplicate samples were assessed for each gene of interest, and GAPDH was used as a control gene. Relative expression levels were determined by the 2^−ΔΔCt^ method. Primer sequences used are given in Table S2.

### Generation of *Dnmt3a* knockout ASCs lines

Guide RNA sequences were cloned into the plasmid px459 (Addgene, 62988). px459 containing *Dnmt3a* gRNAs were co-transfected into digested single ASC by Lipofectamine 2000 (Thermo Fisher Scientific). Single cell derived colonies were picked and expanded individually. Genomic DNA of colonies were extracted using the DNeasy Blood & Tissue Kit, which was further analyzed by genomic PCR. Colonies with the deletion of *Dnmt3a* locus were identified. *Dnmt3a* knockout ASCs (*Dnmt3a^−/-^* ASCs) were cultured in ABCL medium without puromycin. Guide RNA sequences and genotyping primer sequences used are given in Table S2.

### RNA extraction and sequencing

Total RNA were extracted from approximately one million to two million cells using RNeasy Mini Kit (QIAGEN) according to the recommendation of manufacturer and then NEBNext® Poly (A) mRNA Magnetic Isolation Module was used to isolate mRNA from total RNA. Using mRNA as input, the first and second strand cDNAs were synthesized by NEBNext® RNA First Strand Synthesis Module and NEBNext® Ultra II Non-Directional RNA Second Strand Synthesis Module, respectively. Final libraries were prepared using KAPA Hyper Prep Kits (8 PCR cycles) and sequenced on HiSeq4000 platform.

### RNA-seq data analysis

Before alignment, raw data were first trimmed to remove reads with more than 10% low quality bases and to trim adaptors. Then the clean reads were mapped to mouse reference genome (mm10) with Tophat (2.0.12) with default settings (40). HTSeq (0.6.1) was used to do the reads counting, and then RefSeq gene expression level was estimated by RPKM method (Reads per kilobase transcriptome per million reads). Data of RNA-seq of S/L-ESCs and EpiSCs (GSE119985) were downloaded from previous study. Differentially expressed genes (DEGs) in different samples were determined by edgeR package with fold-change ≥ 2 and p value ≤ 0.5 (41). Unsupervised hierarchical clustering (UHC) analysis was performed by the R hclust function. Heatmaps of select genes were performed using R heatmap.2 function. Principal component analysis (PCA) analysis was performed with the R prcomp function. Gene ontology analysis was performed using Metascape (http://metascape.org). Trend analysis of DEGs was performed using Short Time-series Expression Miner (STEM) software (24).

### Genomic DNA isolation and WGBS library preparation

Following the manufacturer‟s instructions, genomic DNA was extracted from stem cells using the DNeasy Blood & Tissue Kit (Qiagen). Remaining RNA was removed by treating with RNase A. Three replicated samples from each of these stem cells were used for library preparation to ensure repeatability of experiment. In short, 2 μg of genomic DNA spiked with 10 ng of lambda DNA were fragmented to about 300 bp with Covaris S220. Next, end repair and A-ligation were performed to the DNA fragments. Methylated Adaptor (NEB) was then ligated to the DNA fragments. In order to reach >99% bisulfite conversion, the adaptor-ligated DNA was treated twice using EZ-96 DNA Methylation-Direct™ MagPrep (Zymo Research). The resulting single-strand DNA fragments were amplified by 4 PCR cycles using the KAPA HiFi HotStart Uracil+ ReadyMix (2×). At last, the libraries were sequenced on HiSeq4000 platform to generate 150-bp paired-end reads.

### DNA methylation analysis

Whole genome bisulfite sequencing reads were trimmed with Trim Galore (v0.3.3) to remove adaptors and low quality bases. Then we used Bismark (v0.7.6) (42) to map the clean reads to mouse reference genome (mm10) with a paired-end and non-directional model, then the unmapped reads were realigned to the same genome with a single-end and non-directional model. PCR duplications were removed with command „samtools rmdup‟ (v0.1.18). WGBS data of 2i/L-ESCs and S/L-ESCs were downloaded from previous study (GSE98517) (8) and identically processed. The global DNA methylation level, estimated using a 2 kb window across the genome, and DNA methylation level in each genomic regions was estimated based on 3x CpG sites (CpGs covered more than 3 times). Only regions with more than 3 CpGs covered were retained. Genomic annotation, like exons, introns and repeat regions were downloaded from UCSC genome browser. Promoters were regions 1 kb upstream and 0.5 kb downstream of transcription start sites (TSS). Imprint control regions (ICR) were obtained from previous study (43), for the low coverage of published S/L-ESCs data, DNA methylation level on ICRs were estimated based on 1x CpG sites. Locations of ICRs were converted with UCSC LiftOver from mm9 to mm10.

### Data availability

These RNA-seq data are available through the NCBI Sequence Read Archive (SRA) under the ID PRJNA601004 (https://dataview.ncbi.nlm.nih.gov/object/PRJNA601004?reviewer=ckal7bagkptogce20v1qf6mp2o). WGBS data have been deposited in the NCBI Gene expression omnibus (GEO) under accession number GSE142799 (https://www.ncbi.nlm.nih.gov/geo/query/acc.cgi?acc=GSE142799). All data that support the conclusions in the study are available from the authors on reasonable request.

### Statistical analysis

All values are depicted as mean ± SD. Statistical parameters including statistical analysis, statistical significance, and n value are reported in the Figure legends and supporting Figure legends. Statistical analyses were performed using Prism Software (GraphPad Prism version 6). The significance of differences was measured by an unpaired two-tailed Student’s *t* test was employed. A value of *p* < 0.05 was considered significant.

## Author contributions

B.W., F.T., M.A.S., X.L. and S.B. designed the experiments. B.W., Y.L., B.Z., Y.W. and Y.F. conducted the experiments; B.L. analysed the RNA-seq data. L.L. and J.G. prepared whole-genome bisulfite sequencing experiment and analyses BS-seq data. C.C. and S.L. helped proof to the manuscript.

## Acknowledgements

We are grateful to Dr Guoliang Xu for the gift of the *Dnmt3l* knockout ESCs. We thank Dr. Juan Li for critical reading of manuscript. Part of the bioinformatics analysis was performed on the Computing Platform of the Peking–Tsinghua Center for Life Science at Peking University.

## Competing interests

The authors declare no competing interest.

## Notes

### Competing Interest Statement

The authors have declared no competing interest.

