## Supplemental Figures and Tables for "DNA methylation is indispensable for leukemia inhibitory factor dependent embryonic stem cells reprogramming"

### SI Appendix information for

### Figure S1

**A**

| Cell line | Original medium | Derived L-ESCs | Repeated times |
|---|---|---|---|
| W1 | 2i/LIF | yes | 1 |
| W2 | 2i/LIF | yes | 3 |
| W4 | 2i/LIF | yes | 2 |
| W5 | 2i/LIF | yes | 2 |
| W6 | 2i/LIF | yes | 3 |
| SQ3.3 | 2i/LIF | yes | 3 |
| Total | 2i/LIF | yes | 16 |

**B**

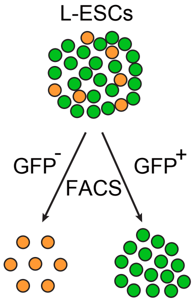


| Cell line | medium | No. Sorting | Passage | GFP+ (%) |
|---|---|---|---|---|
| W2 | LIF | 1 | P23 | 66 |
| | | 2 | P29 | 98 |
| | | 3 | P31 | 99 |
| W6 | LIF | 1 | P14 | 56 |
| | | 2 | P18 | 78 |
| | | 3 | P21 | 91 |
| | | 4 | P33 | 78 |
| | | 5 | P42 | 98 |
| W4 | LIF | 1 | P34 | 97 |
| W5 | LIF | 1 | P34 | 98 |
| W2 | 2i/LIF | 1 | P15 | 98 |

**C**

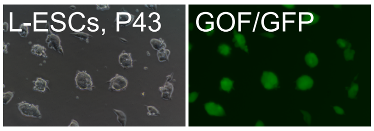


**D**

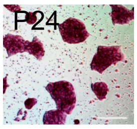


**E**

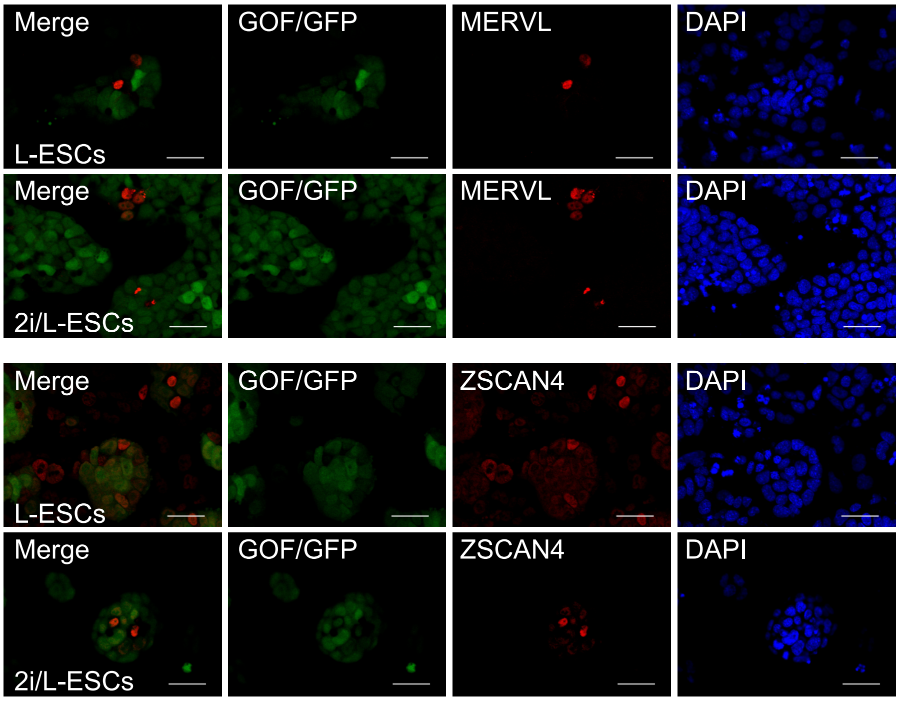


**F**

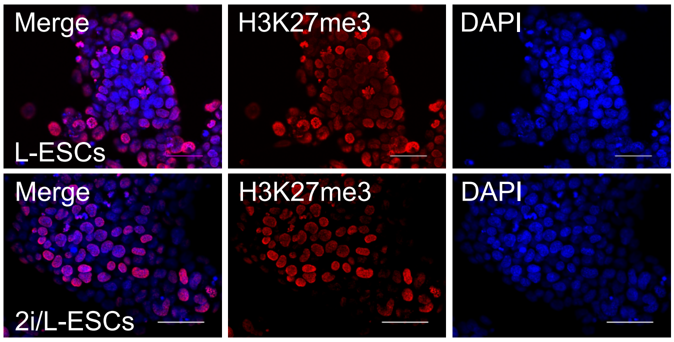

**Figure S1. Characteristics of L-ESCs.**

(A) Derivation of L-ESCs from 2i/L-ESCs.

(B) The summary of fluorescence-activated cell sorting (FACS) based on GOF/GFP positive cells in different passages L-ESCs.

(C) Morphology of L-ESCs at p43.

(D) Alkaline phosphatase (AP) staining on L-ESCs (p24). Scale bars, 100 μm.

(E) Immunostaining of MERVL and ZSCAN4 in 2i/L-ESCs and L-ESCs. Scale bars, 50 μm.

(F) Immunostaining of H3K27me3 in2i/L-ESCs and L-ESCs. Scale bars, 50 μm.

**Figure S2**

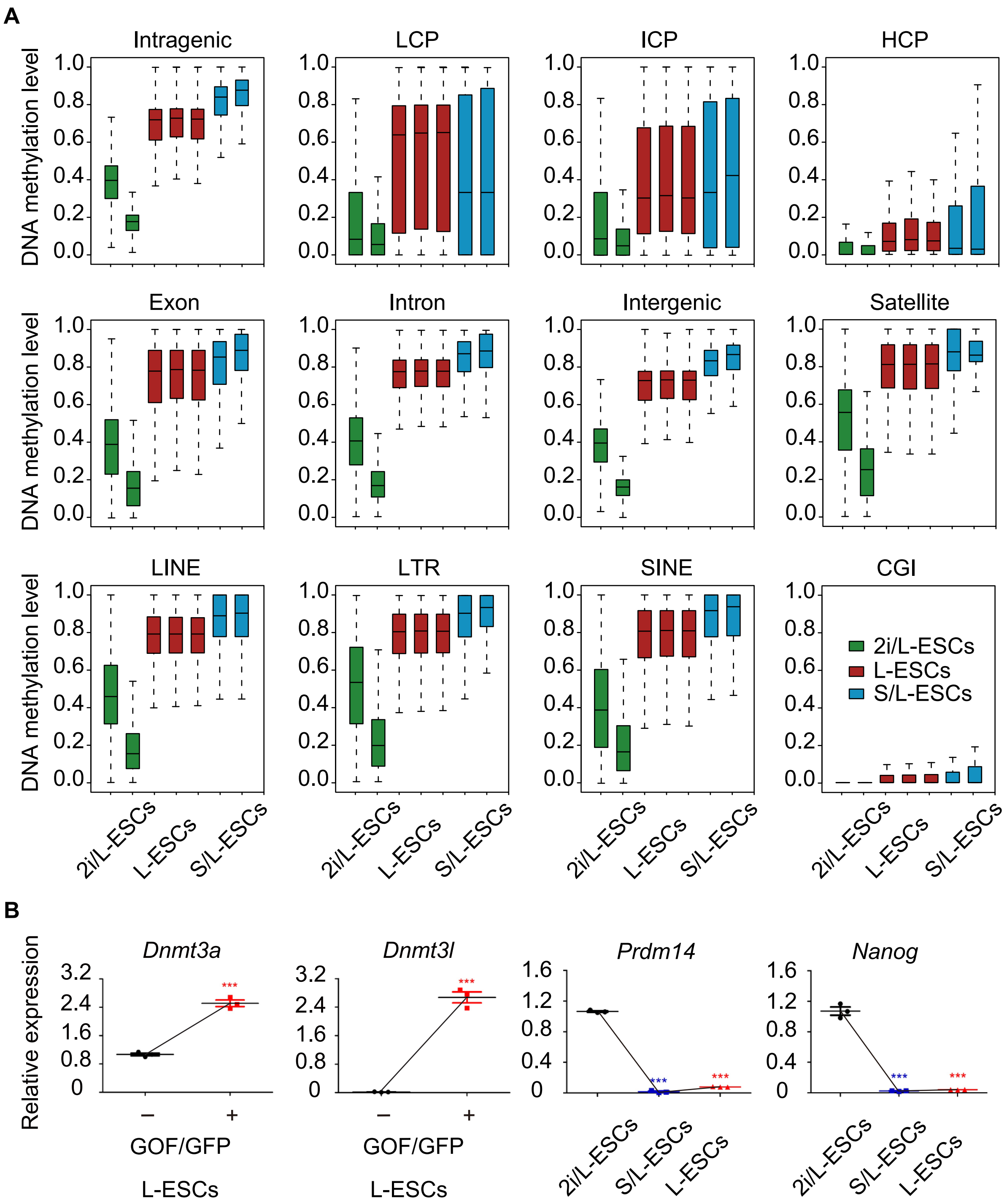
A
Intragenic
LCP
ICP
HCP
Exon
Intron
Intergenic
Satellite
LINE
LTR
SINE
CGI
DNA methylation level
1.0
0.8
0.6
0.4
0.2
0.0
2i/L-ESCs
L-ESCs
S/L-ESCs
B
Dnmt3a
Dnmt3l
Prdm14
Nanog
Relative expression
3.2
2.4
1.6
0.8
0
1.2
0.4
***
–
+
GOF/GFP
L-ESCs
2i/L-ESCs
S/L-ESCs
L-ESCs

**Figure S2. Upregulation of DNA methylation level in L-ESCs.**

(A) DNA methylation level at various genomic features.

(B) Relative expression of *Dnmt3a* and *Dnmt3l* measured by qPCR in GOF/GFP positive and negative L-ESCs; Relative expression of *Prdm14* and *Nanog* measured by qPCR in 2i/L-ESCs, S/L-ESCs and L-ESCs. Error bars are mean ± SD (n = 3). *P* values were calculated by two tailed Student's *t*-test, $p < 0.05$.

**Figure S3**

**A**

| Cell line | Original medium | S/L induction and L-ESCs reprogramming | Repeated times |
|---|---|---|---|
| W2 | 2i/L | yes | 3 |
| W6 | 2i/L | yes | 4 |
| W4 | 2i/L | yes | 4 |
| SQ3.3 | 2i/L | yes | 2 |
| J1 | S/L | yes | 3 |
| *Dnmt3l-/-* | S/L | yes | 3 |

**B**

| Cell line | W5 | | 329 | | | |
|---|---|---|---|---|---|---|
| Culture medium | 2i+LIF | 2i+LIF | ABCL | ABCL | LIF | LIF |
| Number of $AP^+$ colonies | 1673 | 1862 | 3425 | 3672 | 1660 | 1548 |

**C**

| Cell line | Original medium | Reprogrammed L-ESCs | Repeated times |
|---|---|---|---|
| G3 | ABCL | yes | 3 |
| G3-AH | ABCL | yes | 2 |
| G12 | ABCL | yes | 1 |
| 329 | ABCL | yes | 3 |
| *Dnmt3a-/-* | ABCL | yes | 2 |

**Figure S3. Serum improves the efficiency of L-ESCs reprogramming.**

(A) Derivation of L-ESCs from 2i/L-ESCs after 5 days S/L medium culture.

(B) Summary of AP positive cloning numbers on 2i/L-ESCs, ASCs and L-ESCs, when 2,000 cells were seeded into 6-well cell culture plate and 6 days culture.

(C) ASCs can be efficiently reprogrammed into LIF-dependent ESCs.

**Figure S4**

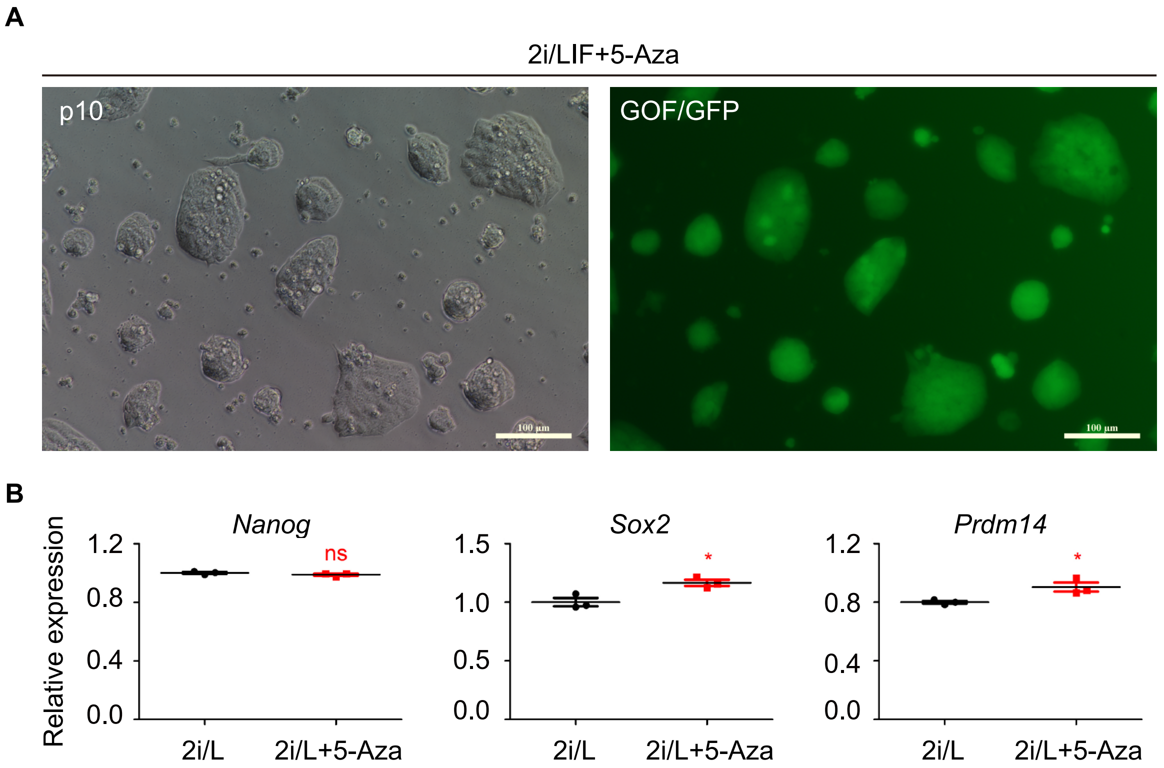

A
2i/LIF+5-Aza
p10
GOF/GFP
B
Nanog
Sox2
Prdm14
Relative expression
1.2
0.8
0.4
0.0
1.5
1.0
0.5
0.0
1.2
0.8
0.4
0.0
ns
*
*
2i/L
2i/L+5-Aza
2i/L
2i/L+5-Aza
2i/L
2i/L+5-Aza

**Figure S4. DNA methylation was indispensable for L-ESCs self-renew.**

(A) 2i/L-ESCs were treated with 5-Aza after p10, 2i/L-ESCs retained typical dome-shaped clonal morphology. Scale bars, 100 μm.

(B) Relative expression of *Nanog*, *Sox2* and *Prdm14* measured by qPCR in 2i/L-ESCs after 3 days 5-Aza treatment. Error bars are mean ± SD (n = 3). *P* values were calculated by two tailed Student's *t*-test, $p < 0.05$.

**Figure S5**

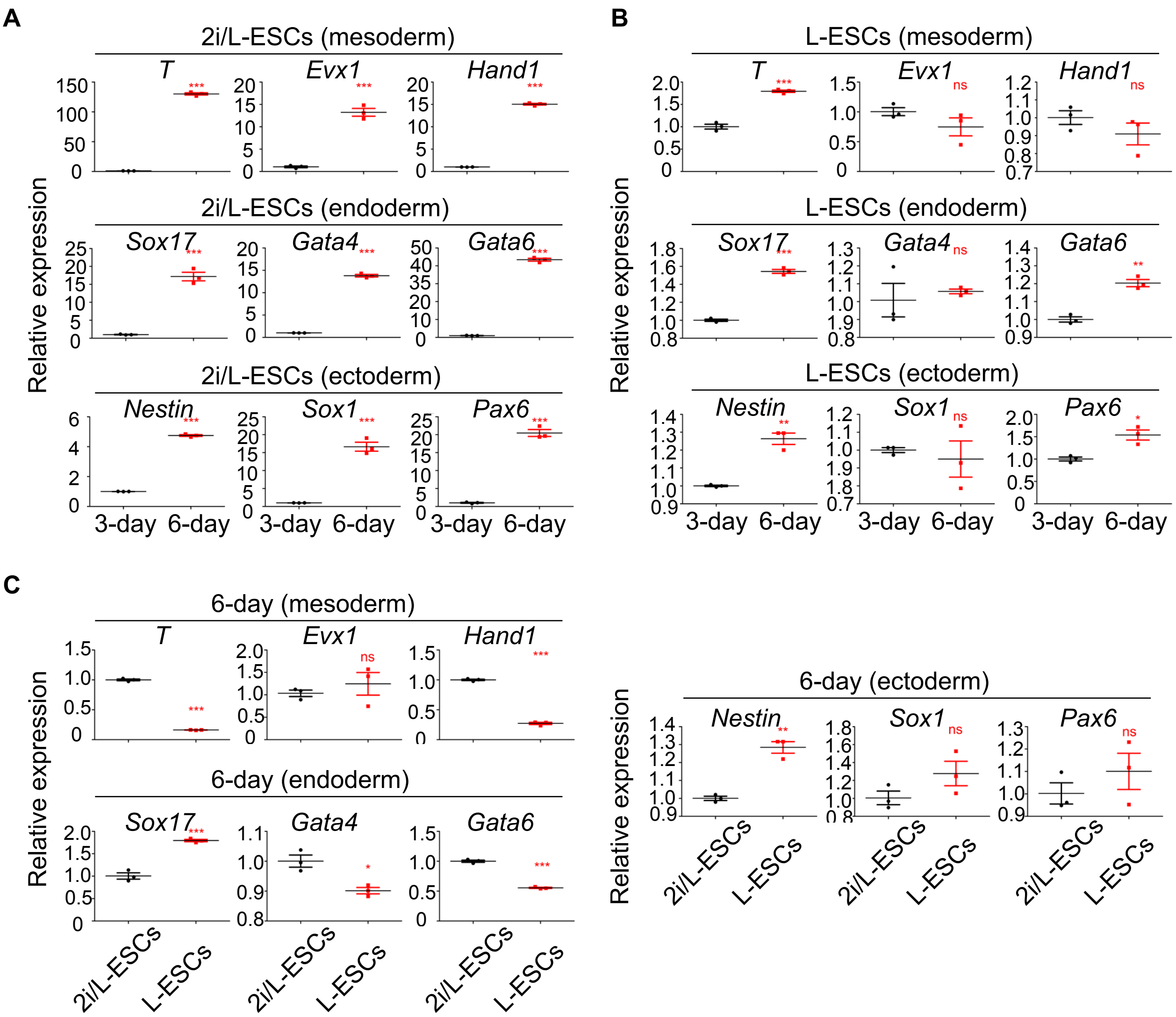
A
2i/L-ESCs (mesoderm)
T
Evx1
Hand1
2i/L-ESCs (endoderm)
Sox17
Gata4
Gata6
2i/L-ESCs (ectoderm)
Nestin
Sox1
Pax6
Relative expression
3-day
6-day
B
L-ESCs (mesoderm)
L-ESCs (endoderm)
L-ESCs (ectoderm)
ns
C
6-day (mesoderm)
6-day (endoderm)
6-day (ectoderm)
2i/L-ESCs
L-ESCs

**Figure S5. The pluripotency of L-ESCs *in vivo* and *in vitro*.**

(A) Relative expression of mesoderm, endoderm and ectoderm genes measured by qPCR, after 2i/L-ESCs were 3 days and 6 days *in vitro* differentiation. Error bars are mean ± SD (n = 3). *P* values were calculated by two tailed Student's *t*-test, $p < 0.05$.

(B) Relative expression of mesoderm, endoderm and ectoderm genes measured by qPCR, after L-ESCs were 3 days and 6 days *in vitro* differentiation. Error bars are mean ± SD (n = 3). *P* values were calculated by two tailed Student's *t*-test, $p < 0.05$.

(C) Relative expression of mesoderm, endoderm and ectoderm genes measured by qPCR, after 2i/L-ESCs and L-ESCs were 6 days *in vitro* differentiation. Error bars are mean ± SD (n = 3). *P* values were calculated by two tailed Student's t-test, p < 0.05.

**Table S1. WGBS data coverage and conversion efficiency (Related to Fig 3A)**

| Sample | Total sequenced bases(Gb) | Total sequenced reads | Reads after trimming | Unique mapped reads | Mapping efficiency | Coveraged bases (1x) |
|---|---|---|---|---|---|---|
| L-ESCs_rep1 | 23.36 | 155,760,956 | 151,382,966 | 98,603,675 | 65.14% | 2,317,989,827 |
| L-ESCs_rep2 | 26.20 | 174,654,234 | 168,555,054 | 106,718,870 | 63.31% | 2,334,743,972 |
| L-ESCs_rep3 | 24.60 | 163,969,284 | 158,835,308 | 103,120,363 | 64.92% | 2,346,240,160 |

| Sample | Fraction of genome covered | Bisulfite Conversion Rate | No.of unique CpG covered (1x) | No.of unique CpG covered (3x) | No.of unique CpG covered (5x) |
|---|---|---|---|---|---|
| L-ESCs_rep1 | 85.05% | 99.96% | 30,757,370 | 14,829,901 | 5,277,450 |
| L-ESCs_rep2 | 85.66% | 99.96% | 30,976,871 | 15,454,751 | 5,806,380 |
| L-ESCs_rep3 | 86.08% | 99.96% | 31,620,932 | 15,560,785 | 5,654,681 |

**Table S2. RT-qPCR Primers and Guide RNA sequences**

| Gene name | Forward Primer | Reverse Primer |
|---|---|---|
| *Nanog* | CTTTCACCTATTAAGGTGCTTGC | TGGCATCGGTTCATCATGGTAC |
| *Prdm14* | CCTGAACAAGCACATGAGA | TGCACTTGAAGGGCTTCTCT |
| *Gata4* | TTCCTCTCCCAGGAACATCAAA | GCTGCACAACTGGGCTCTACTT |
| *Gata6* | TGCTGGAAATTGCAACAAACC | GTCACGTGGTACAGGCGTCA |
| *Sox17* | GTCAACGCCTTCCAAGACTTG | GTAAAGGTGAAAGGCGAGGTG |
| *Brachyury* | GAACCTCGGATTCACATCGT | TTCTTTGGCATCAAGGAAGG |
| *Evx1* | CCAGTGACCAGATGCGCCGATAC | TCCTTCATGCGCCGGTTCT |
| *Hand1* | TCAAAAAGACGGATGGTGGT | GCGCCCTTTAATCCTCTTCT |
| *Dnmt3a* | GACTCGCGTGCAATAACCTTAG | GGTCACTTTCCCTCACTCTGG |
| *Dnmt3l* | CGGAGCATTGAAGACATC | CATCATCATACAGGAAGAGG |
| *Sox2* | GCGGCGGAAAACCAAGA | CCGGGAAGCGTGTACTTATCC |
| *Nestin* | CTCGAGCAGGAAGTGGTAGG | TTGGGACCAGGGACTGTTAG |
| *Sox1* | GGCCGAGTGGAAGGTCATGT | TCCGGGTGTTCCTTCATGTG |
| *Pax6* | GCAGATGCAAAAGTCCAGGTG | CAGGTTGCGAAGAACTCTGTTT |
| *GAPDH* | ATGGTGAAGGTCGGTGTGAAC | TCGCTCCTGGAAGATGGTGATG |

**Guide RNA sequences**

| | |
|---|---|
| *Dnmt3a* sgRNA1 | CACCGCTCATACTCAGGCTCATCGT |
| *Dnmt3a* sgRNA1-CS | AAACACGATGAGCCTGAGTATGAGC |
| *Dnmt3a* sgRNA2 | CACCGGACCCTGCTTCTCCGACTG |
| *Dnmt3a* sgRNA2-CS | AAACCAGTCGGAGAAGCAGGGTCC |

**Genotyping primer**

| | |
|---|---|
| *Dnmt3a* 89925 Forward Primer | GCCTTGGCTGTGTGAGATTTG |
| *Dnmt3a* 90586 Reverse Primer | ATCCTGGAGCCCCAAAGAGC |
